# Relationship between maximal incremental and high-intensity interval exercise performance in elite athletes

**DOI:** 10.1101/856237

**Authors:** Shih-Chieh Chang, Alessandra Adami, Hsin-Chin Lin, Yin-Chou Lin, Carl P.C. Chen, Tieh-Cheng Fu, Chih-Chin Hsu, Shu-Chun Huang

**Affiliations:** Department of Physical Medicine & Rehabilitation, Chang Gung Memorial Hospital, Linkou, Taiwan; Department of Kinesiology, University of Rhode Island, Kingston, RI, USA; Department of Physical Medicine & Rehabilitation, Chang Gung Memorial Hospital, Taoyuan branch, Taiwan; College of Medicine, Chang Gung University, Kwei-Shan, Tao-Yuan County, Taiwan; Department of Physical Medicine and Rehabilitation, Chang Gung Memorial Hospital, Keelung, Taiwan; Healthy Aging Research Center, Chang Gung University, Taoyuan City, Taiwan

**Keywords:** exercise testing, oxygen uptake, heart rate, near-infrared spectroscopy, HIIT

## Abstract

It remains unclear whether the number of total bouts to limitation (B_lim_) in high-intensity interval testing (HIIT) differs among individuals, no matter if performed at the same relative intensity. This study aimed to explore the physiologic factors determining tolerance to effort during a HIIT. Forty-seven female participants (15-28 years old) were included: 23 athletes from Taiwan national or national reserve teams, and 24 moderately-active female. Each participant underwent maximal incremental (INC; modified-Bruce protocol) cardiopulmonary exercise testing and HIIT on treadmill, on separate days. HIIT protocol alternated a 1-min effort at 120% of the maximal speed and the same slope reached at the end of INC, with a 1-min rest, until volitional exhaustion. Gas-exchanges, and muscle oxygenation at right vastus lateralis by near-infrared spectroscopy, were continuously recorded. Additionally, bioelectrical impedance was utilized for body composition analysis. The result showed that B_lim_ differed greatly (range: 2.6 to 12) among participants. Stepwise regression revealed that B_lim_ was determined primarily by oxygen consumption (VO_2_) and heart rate (HR) at second-minute recovery; and, muscle tissue saturation index at peak of INC (R=0.644). Also, age and percent body fat were linearly correlated with B_lim_ (adjusted R=−0.475, −0.371, p<0.05). Therefore, HIIT performance is determined by fast recovery of VO_2_ and HR, rather than maximal VO_2_ or muscle oxygenation recovery. Moreover, capacity to sustain a HIIT declines with age since as early as late adolescent. Further investigations on which factors should be manipulated to further improve athletes performance are warrant.

## INTRODUCTION

Incremental exercise testing (INC) is nowadays widely used to assess cardiopulmonary fitness among various populations, from élite athletes to semi-professional players, to chronic cardiovascular and lung disease patients [1, 2]. The INC provides quantification of whole-body all-out performance and, in the athletic world, it has become the gold-standard evaluation to identify exercise intensity zones upon which athletic training programs are designed.

In many competitive ball games, such soccer, basketball, rugby, badminton, a typical field performance is interspersed with multiple intervals (i.e. combination of succeeding effort and recovery phases in series). At high professional level, the intensity of this field performance can be compared to a high-intensity interval workout. This latter has been for decades a preferred athletic training method to increase not only whole-body aerobic capacity, but also to manipulate the response of the peripheral skeletal muscle system in reducing the time of activation and recovery kinetics of oxygen transport metabolism [3] as to, consequently, reduce the delay between mechanical request (i.e. exercise task) and muscle metabolic response (the so called metabolic phase or Phase II [4]. It is likely that the performance in high-intensity interval testing (HIIT) is more related to the court performance of athletes competing in the above-mentioned team sports. If that is true, the traditional INC should not be the first testing choice to evaluate ball games players performance and to base upon the seasonal training schedule. Nonetheless, HIIT has seldomly been used as testing protocol [5, 6] and INC results are still used to planning the training calendar, which is largely a combination of high-intensity exercise workloads and interval training phases.

In 2017, a group of reserve athletes from Taiwan national soccer, basketball and badminton teams came to our laboratory at the Chang Gung Memorial Hospital for a series of performance routine evaluations. In that occasion we administered a traditional INC protocol and a newly designed HIIT. Similar to the concept of “time to limitation” (i.e. T_lim_) in constant work rate (CWR) testing, we applied the same approach to HIIT test, using the number of bouts to limitation (B_lim_) and a supra-maximal intensity exercise (i.e. 120% INC maximal velocity) as indexes to assess the overall cardiopulmonary fitness in ball games sport players. Repeated transitions between supra-maximal intensity and recovery phase involve anaerobic and aerobic energy metabolisms both during and immediately after the exercise.

The recovery capacity is an important physiologic determinant affecting the performance in many types of ball games but is not well assessed with the traditional INC or CWR testing. In the same relative intensity during CWR, the time to limitation only slightly varies from person to person based on the concept of power duration hyperbolic curve (for example, T_lim_ is 6 ± 2 minutes at 80% peak work rate in every subject) [7]. In contrast, in HIIT, whether B_lim_ varies considerably or not among individuals is still unclear.

Therefore, the main aim of this descriptive study was to determine which factors are most strongly associated with the bouts to limitation in HIIT. We hypothesized that B_lim_ in HIIT, unlike T_lim_ in CWR testing, has a wide discrepancy among individuals given the same relative intensity and exercise-recovery duration and pattern, which we suggest is primarily due to individual recovery capacities.

## MATERIALS AND METHODS

### Participants

This study enrolled 47 athletes and moderately active (MA) female participants. The athletes were reserves for the national Taiwan soccer, basketball and badminton teams. The participants assigned to the MA group were young women participating in moderate intensity exercise for at least 60 min in total weekly [8]. The experiment protocol was approved by the Chang Gung Memorial Hospital Institutional Review Board. All participants provided a written informed consent after receiving oral and written explanations of the experimental procedures and associated risks. This research was performed in accordance with the ethical standards of the Declaration of Helsinki.

### Protocol

Participants came twice at our laboratories at the Chang Gung Memorial Hospital to perform an INC maximal test and a HIIT. Each subject was instructed to refrain from vigorous exercise or caffeine intake for the 24 h prior testing, and to have at least 8-hour sleep the night before the test. All assessments took place approximately at the same time of the day under controlled environmental conditions (24°C, 63% humidity).

#### Anthropometric and Body Composition Evaluation

At the beginning of the visit, basic anthropometric characteristics (height and weight) were taken. Afterwards, whole-body composition was determine using the InBody s10 (Seoul, Korea) and by measuring the electrical resistance to four different frequencies (5, 50, 250, and 500 kHz) [9-11]. Each participant lied on a padded table for the entire duration of the testing, and sensors to measure electrical resistance were placed at the level of each body segment following the manufacturer instructions. To undergo this 20-min procedure, participants were instructed to fast for 2 hours prior to the test.

#### Cardiopulmonary testing

During the first visit a maximal INC test was carried out. The test started with 1 min of walking at 1 mile/h, followed by an incremental modified-Bruce protocol conducted until volitional exhaustion was reached. Immediately after exhaustion, an active recovery phase at individual walking pace was administered for 1 min, followed by 3 min of passive recovery. The INC was defined as maximal when the following criteria were present: (i) a plateau in VO_2_ between the final 2 stages, (ii) HR exceeded 85% of its predicted maximum, and (iii) respiratory exchange ratio exceeded 1.15 [2].

During the second visit, that took place at least 24 hours after the first one, a supra-maximal HIIT was performed. The speed was set at 120% of the maximal velocity reached during the INC, and the slope was the same as the INC final stage. The HIIT protocol consisted of intermittent 1-min sprinting interspersed with 1-min passive recovery, until exhaustion. Total number of bouts completed before exhaustion was recorded (B_lim_).

### Measurements

All exercise assessments were performed on an electromechanically braked treadmill (VIASYS™), while minute ventilation (V_E_), VO_2_, and carbonic dioxide production (VCO_2_) were measured using a computer-based system (MasterScreen CPX, Cardinal-health, Germany). Before each test, the gas analyzers and the turbine flow meter of the system were calibrated following the manufacturer’s instructions and by using a gas mixture of known concentration (FO_2_: 0.16; FCO_2_: 0.05; N_2_ as balance) and an automatically-pumping high and low flow system. Heart rate was determined from the R-R interval on a 12-lead electrocardiogram.

Muscle oxygenation was evaluated by means of a Bluetooth, portable, continuous-wave and spatially-resolved near-infrared spectroscopy (NIRS) (PortaMon, Artinis, The Netherlands). Relative concentrations of deoxy-hemoglobin+myoglobin (HHb), oxy-hemoglobin+myoglobin (O_2_Hb), and tissue saturation index (TSI, %) [12] were continuously recorded during the exercises at the level of the peripheral muscle tissue, ∼1.5 cm beneath the probe (interoptode distance 3 cm). From these measurements, the relative changes in total hemoglobin and myoglobin (THb = HHb + O_2_Hb) were calculated. The NIRS probe was wrapped in a plastic foil and placed longitudinally to the muscle *vastus lateralis* belly on the right thigh, 15 cm above the upper margin of the patella, and secured with an elastic bandage to minimize the possibility that external light influenced the signal.

### Data Analysis

Maximal VO_2_ (VO_2max_) and maximal heart rate (HR_max_) were defined as the highest values obtained during the INC test. For the recovery phase, a three breaths moving average was applied to calculate VO_2_ and HR off-kinetics. Similarly, for the NIRS data (i.e. TSI and O_2_Hb), five 1-s data were used in the process of moving average.

VO_2,_ HR, O_2_Hb, and TSI changes at 0.5, 1, and 2 minutes recovery after INC were calculated as suggested by Turner et al.[13] (Figure 1):

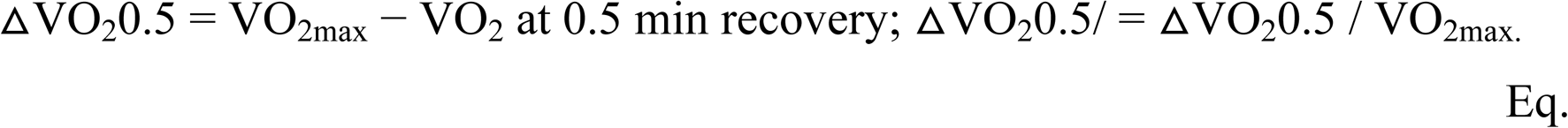

**Figure 1.**
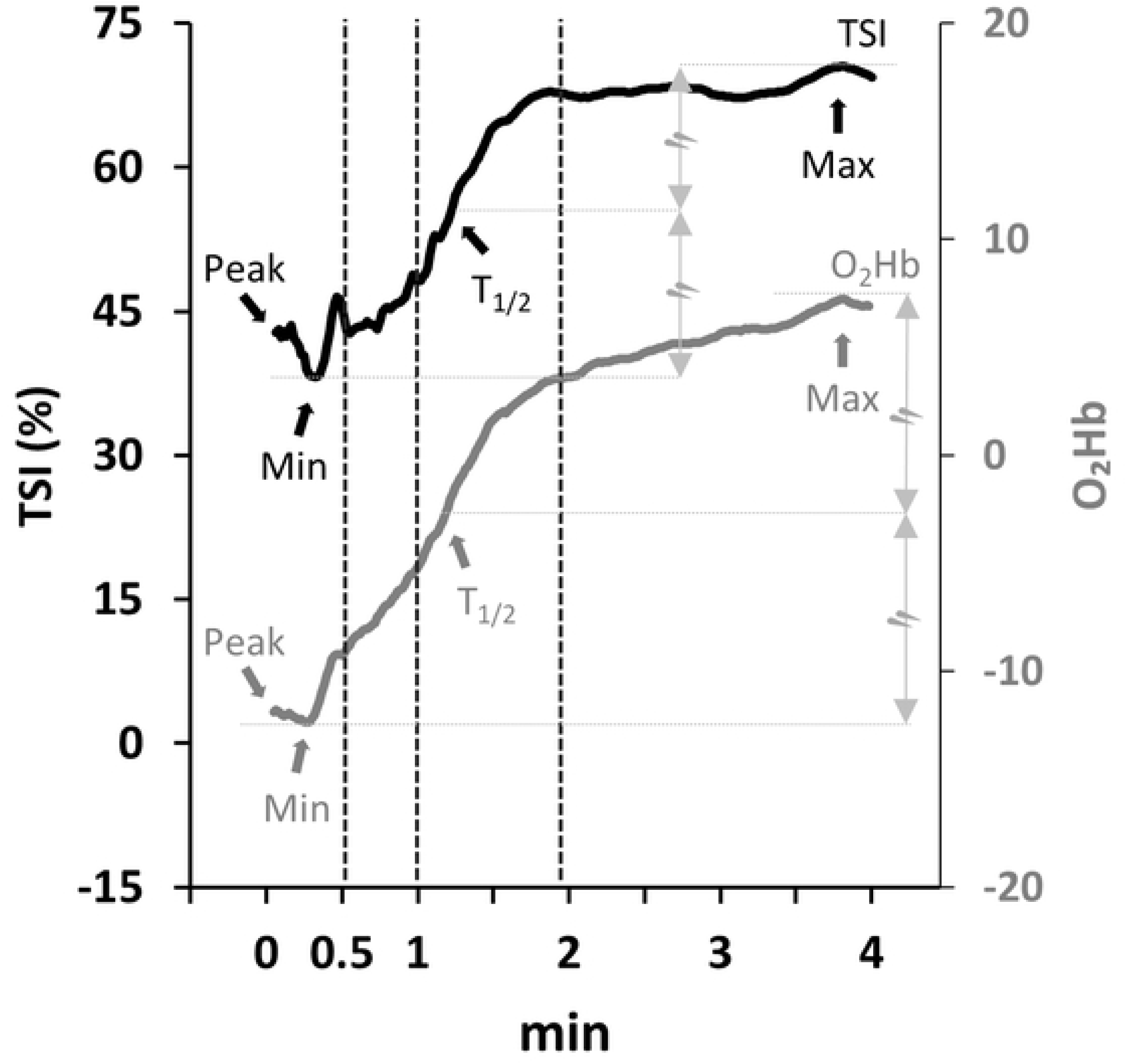
The data of TSI and O_2_Hb after moving average in the recovery phase of INC are graphed.TSI is labeled as black line and O_2_Hb is labeled as grey. Maximal and minimal value of TSI and O_2_Hb are labeled as Max, and Min, respectively. Peak value is the value at peak during INC. T_1/2_ is the time span the variable changes in half. For simplicity, data during exercise is not shown. The calculation formula were as follows. ΔTSI0.5 = TSI at 0.5 min during INC recovery – peak TSI; ΔTSI0.5/ = ΔTSI0.5/ (maximal TSI – minimal TSI) during recovery. The same formula applies to the 1st and 2nd minute, and also O_2_Hb. O_2_Hb: oxy-(hemoglobin+myoglobin), TSI: tissue saturation index, INC: Incremental exercise testing,

Above equation was applied to determine the 1^st^ and 2^nd^ minute change, and change ratio at the same three time points. The same approach was used for HR determination.

In addition, TSIrt_1/2_, O_2_Hbrt_1/2_, VO_2_rt_1/2,_ and HRrt_1/2_ are the time span the value recovers to half.

### Statistical analysis

Data are given as mean and standard deviation (SD). The criterion for significance was set at *p*<0.05. Pearson correlation, correlation matrix and partial correlation were used to determine the degree of association between physiological variables measured during the cardiopulmonary testing and B_lim_. Since the population sample size allocate to final analyses was of 47, the first five parameters with the greatest correlation coefficient were included in the regression model. Forward stepwise linear regression was run to seek for predictors of B_lim_. Analyses were done using SPSS 22.0 (SPSS, Inc., Chicago, IL, USA).

## RESULTS

Forty-seven participants (n=23 athletes; n=24 MA) successfully completed the experimental protocol phases. B_lim_ differed greatly among subjects, ranging from 2.6 to 12. The representative results on the strongest physical parameters correlated to B_lim_ are shown in Figure 2. Among the anthropometric variables, age was significantly moderately correlated to B_lim_ (R=-0.475; *p*=0.008) after adjusting variables with co-linearity, including group, percent body fat (PBF), ΔVO_2_2, △HR2/ and TSI_peak_ (Figure 2A). As to the variables derived from body composition analysis, PBF was significantly associated with B_lim_ (Figure 2B).

**Figure 2.**
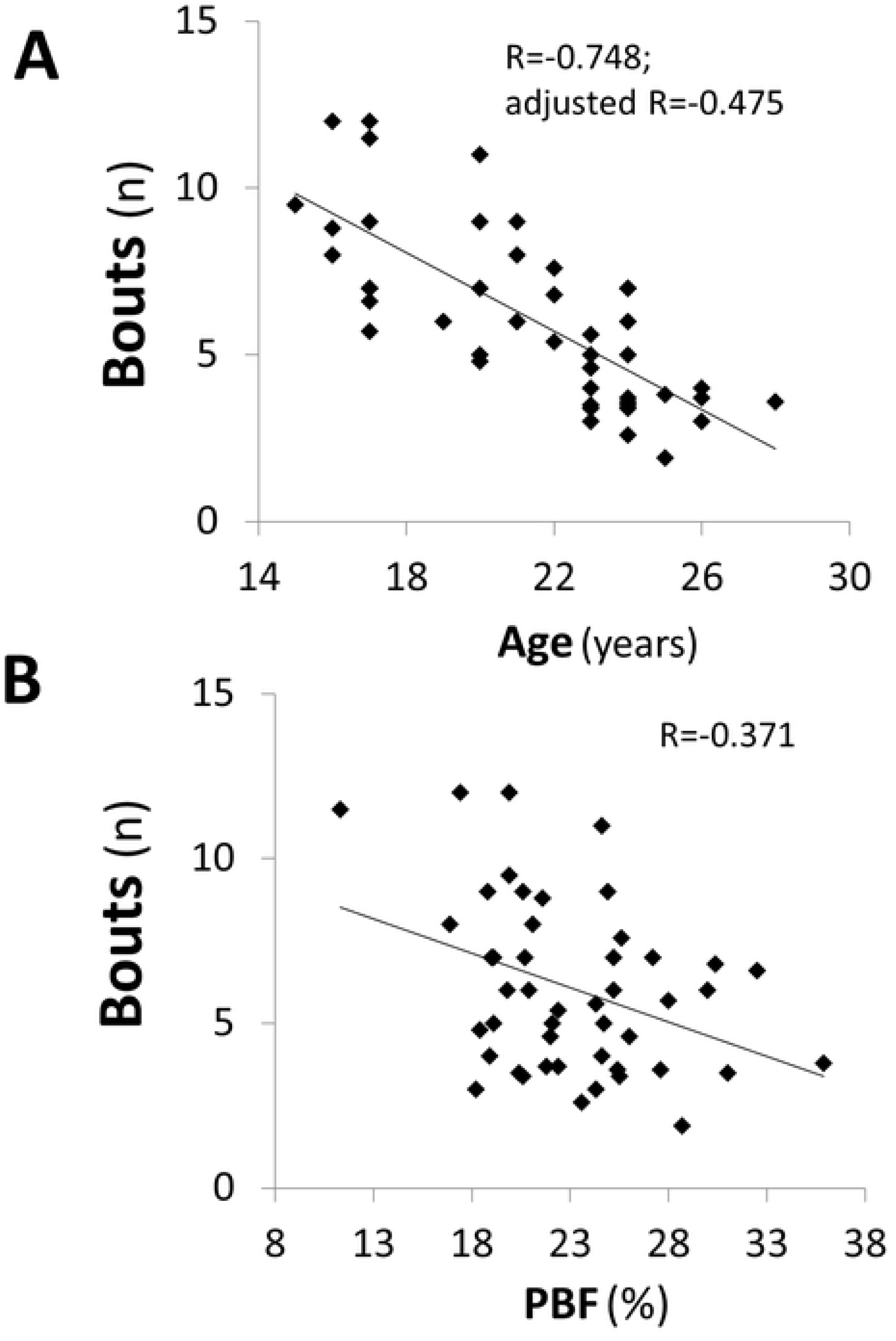
Scatter plots of age and body composition against total number of bouts in high-intensity interval test. In Figure 2A, partial correlation is utilized to adjust the variables with co-linearity, including group, PBF, ΔVO_2_2, △HR2/ and TSI_peak_ PBF, percent body fat; △VO_2_2 = VO_2max_ – VO_2_ at 2 min during recovery; △HR2/ = (HR_max_ – HR at 2 min during recovery) / maximal HR; TSI_peak_, nadir of tissue saturation index during INC

When the HIIT and INC were compared by univariate analyses, several physiological parameters during INC were significantly correlated with B_lim_. Those parameters with a correlation coefficient >0.3 and P<0.05 are shown in S1. The five physical parameters with the highest correlation coefficients (ΔVO_2_2, VO_2max_, △HR2/, TSI_peak_ and △HR2) were put into the forward linear stepwise regression model (Table 1). The explanatory power in model 3 of the multiple linear regression model was 0.415. It revealed that B_lim_ were majorly determined by ΔVO_2_2, △HR2/ and TSI_peak_. Their scatter plots to B_lim_ are shown in Figure 3. Correlations were positive in ΔVO_2_2, △ HR2/, and negative in TSI_peak_ (Figure 3A, 3B and 3C).

**Table 1.**
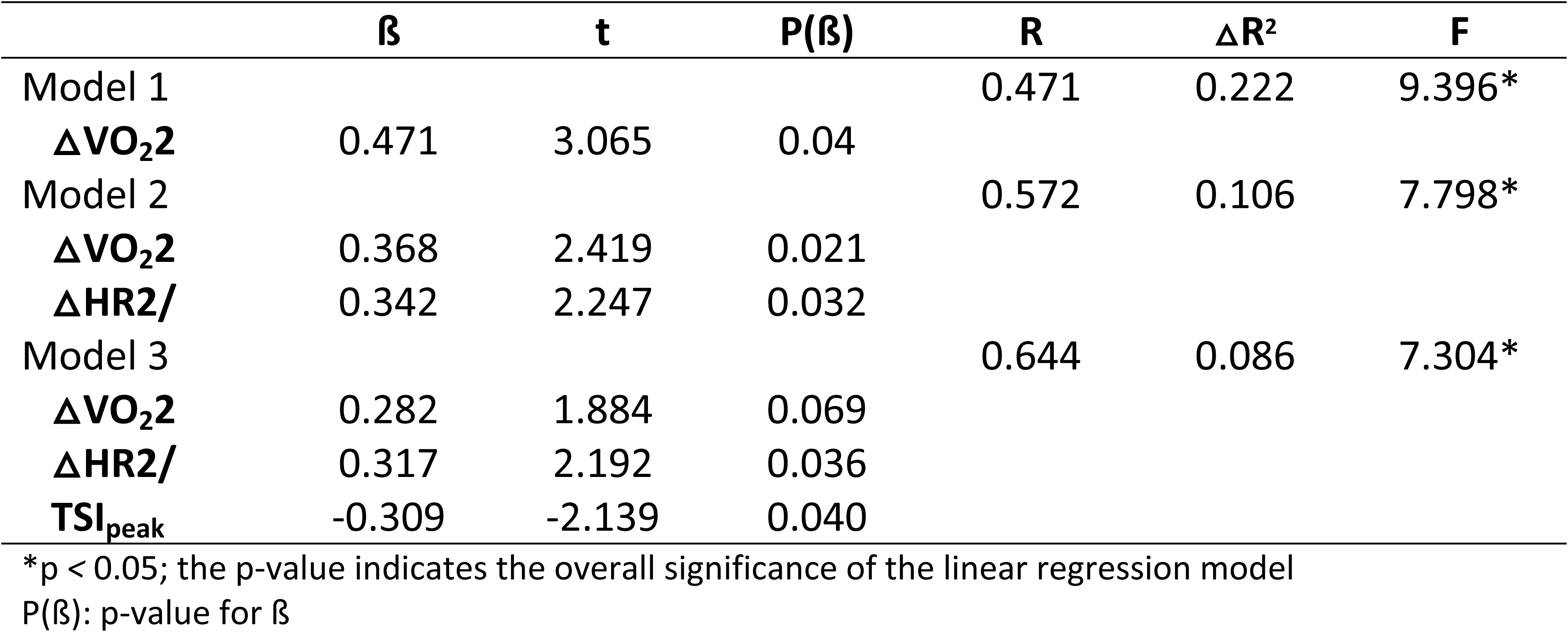
Stepwise linear regression of B_lim_ based on physiologic parameters.

**Figure 3.**
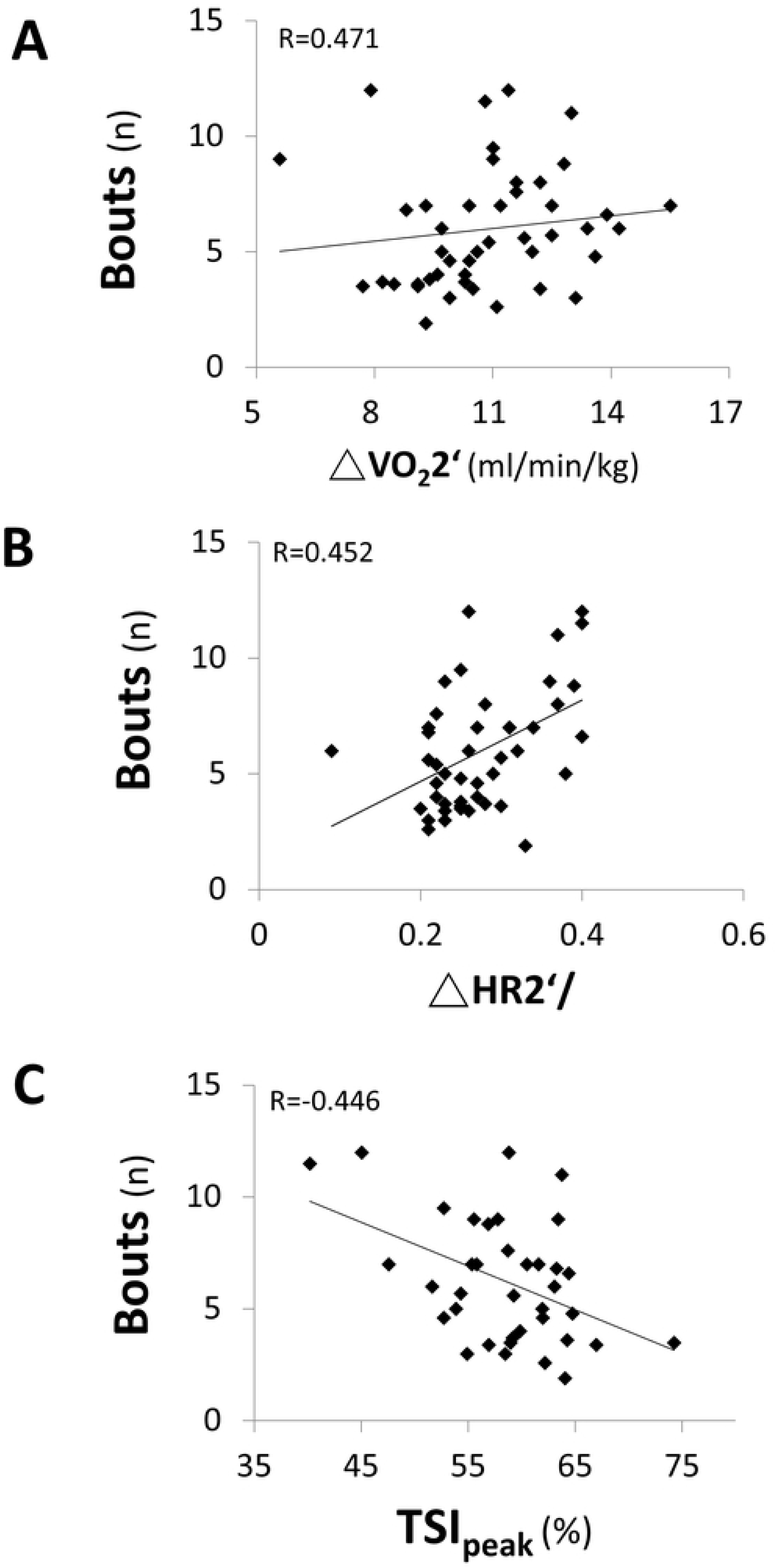
Scatter plots of physical parameters derived from the incremental exercise testing against total number of bouts to limitation in the HIIT △VO_2_2 = VO_2max_ – VO_2_ at 2 min during recovery; △HR2/ = (HR_max_ – HR at 2 min during recovery) / maximal HR; TSI_peak_, nadir of tissue saturation index during INC.

Figure 4 shows an example of a comparison between two subjects with similar VO_2_max but significantly different B_lim_ (12 vs. 7.7). The subjects who tolerated more bouts in HIIT had a faster recovery in VO_2_ and HR and a lower TSI_peak_ value in INC.

**Figure 4.**
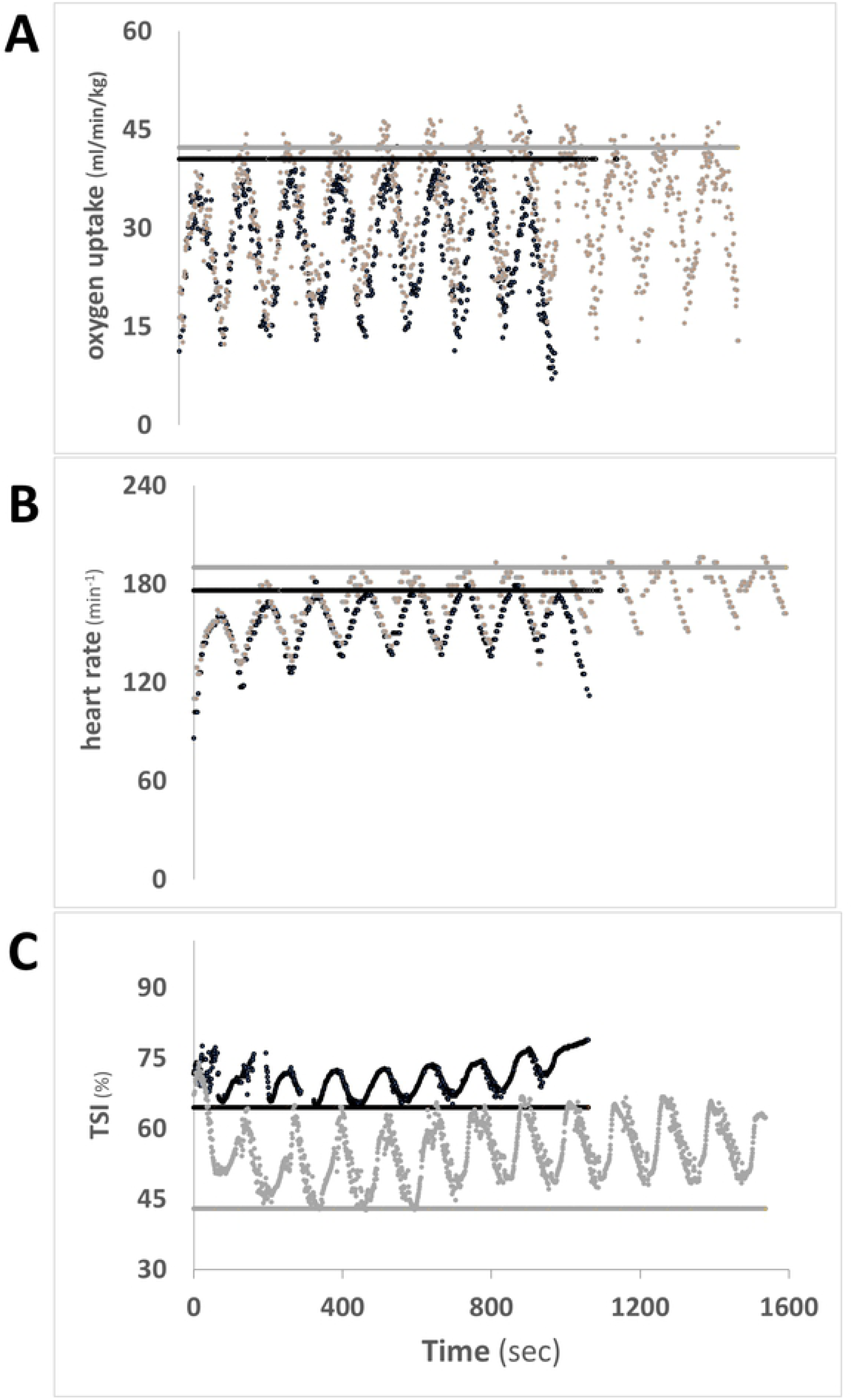
The two subjects had very close maximal VO_2_ in INC but differed greatly in B_lim_(12 vs. 7.7 bouts) in HIIT. The dark and light horizontal lines denote the peak values in their INCs. The subject who tolerated more bouts in HIIT had a faster recovery in VO_2_, HR, and lower TSI_peak_. HIIT, high-intensity interval testing; INC, maximal incremental exercise test; VO_2_, oxygen consumption; TSI_peak_, nadir of tissue saturation index during INC.

The main findings for INC and HIIT are provided in Table 2. As expected, the athletes had a significantly higher VO_2max_, maximal O_2_ pulse, peak V_E_ and lower TSI_peak_ in INC, and better performance in HIIT (B_lim_: 7.8 ± 2.3 vs. 4.2 ± 1.4, *p*<0.05) than MA.

**Table 2.** Physical parameters during INC and HIIT.

|  | Ath | MA |
| --- | --- | --- |
| **INC** |  |  |
| VO<sub>2</sub>max (L.min<sup>-1</sup>.Kg<sup>-1</sup>) | 45.7 ± 6.0* | 37.8 ± 7.6 |
| peak HR (min<sup>-1</sup>) | 185.0 ± 8.4 | 187.4 ± 10.0 |
| peak O<sub>2</sub> pulse (ml) | 12.8 ± 1.8* | 10.4 ± 1.9 |
| EqCO<sub>2</sub> nadir | 24.3 ± 2.2 | 24.1 ± 2.2 |
| EqO<sub>2</sub> nadir | 20.4 ± 2.1 | 21.1 ± 2.7 |
| peak V<sub>E</sub> (L.min<sup>-1</sup>) | 83.0 ± 13.5* | 76.3 ± 14.0 |
| V<sub>E</sub>/VCO<sub>2</sub> slope | 26.8 ± 3.3 | 25.4 ± 6.9 |
| TSI<sub>peak</sub> (%) | 56.4 ± 6.6* | 61.0 ± 5.1 |
| HHb<sub>peak</sub> (μM) | 0.9 ± 5.0 | 2.1 ± 5.0 |
| O<sub>2</sub>Hb<sub>peak</sub> (μM) | -4.1 ± 4.7 | -6.6 ± 4.5 |
| THb<sub>peak</sub> (μM) | -3.3 ± 6.5 | -4.5 ± 7.9 |
| **HIIT** |  |  |
| B<sub>lim</sub> (times) | 7.8 ± 2.3* | 4.2 ± 1.4 |

Value are shown as mean ± SD, \* $p < 0.05$ in Mann-Whitney U test; Ath: athletes; MA: moderately-active subjects; HR: heart rate; V<sub>E</sub>: ventilation; TSI: tissue saturation index; HHb: deoxyhemoglobin; O<sub>2</sub>Hb: oxyhemoglobin; THb: total hemoglobin; TSI<sub>peak</sub>: nadir of tissue saturation index during INC; EqCO<sub>2</sub>: ventilator equivalent for CO<sub>2</sub>; EqO<sub>2</sub>: ventilatory equivalent for O<sub>2</sub>

Main results in the recovery phase of INC are presented in Table 3. HR recovery at the 2_nd_ minute and VO_2_ recovery are faster in Ath than MA. Muscle oxygenation recovery showed no difference between Ath and MA. It is noteworthy that ΔHR2/ and △VO_2_2 that were the major determinants of B_lim_ picked up by the stepwise regression are significantly higher in Ath than MA.

**Table 3.**
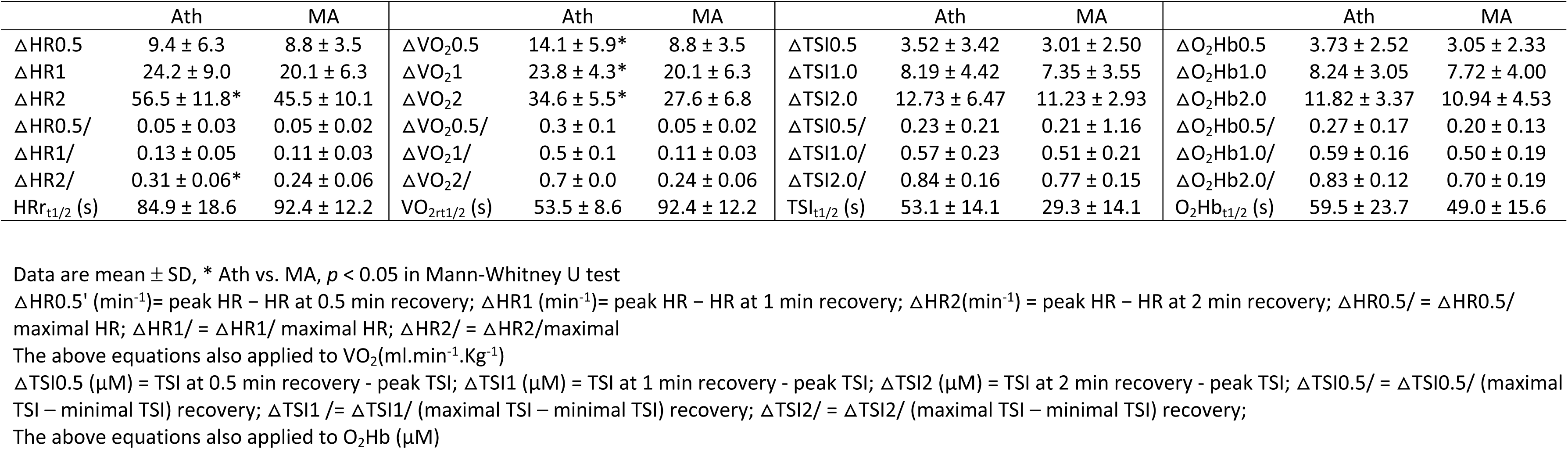
Physiologic response during the recovery phase at the end of the incremental exercise test.

Anthropometric data showed no difference in body weight, body height and BMI except age (19 ± 3 vs. 24 ± 2 years) between Ath and MA. Table 4 shows the body composition data. Ath has higher SMM, SLM, FFM, SMRA, SMTR, SMRL, protein, BCM and lower PBF. The P-value of SMLA and SMLL are 0.051 and 0.055.

**Table 4.** Body composition data.

|  | **Ath** | **MA** |
| --- | --- | --- |
| SMM (Kg) | 23.6 ± 2.3* | 22.2 ± 2.3 |
| Fat (Kg) | 11.8 ± 2.7 | 13.7 ± 3.8 |
| PBF (%) | 21.5 ± 4.1* | 24.9 ± 4.5 |
| SLM (Kg) | 40.4 ± 3.7* | 38.3 ± 3.7 |
| FFM (Kg) | 42.9 ± 3.9* | 40.6 ± 3.9 |
| SMRA (Kg) | 2.1 ± 0.3* | 1.9 ± 0.3 |
| SMLA (Kg) | 2.0 ± 0.3 | 1.9 ± 0.3 |
| SMTR (Kg) | 18.8 ± 1.8* | 17.9 ± 1.7 |
| SMRL (Kg) | 6.9 ± 0.8* | 6.5 ± 0.8 |
| SMLL (Kg) | 6.9 ± 0.7 | 6.6 ± 0.8 |
| Protein (Kg) | 8.5 ± 0.8* | 8.0 ± 0.8 |
| BCM (Kg) | 28.0 ± 2.6* | 26.6 ± 2.6 |
| TBM/FFM (%) | 73.3 ± 0.2 | 73.3 ± 0.2 |

Value are shown as Mean ± SD, \* $p < 0.05$ in Mann-Whitney U test; Ath: athletes; MA: moderately-active subjects; SMM: Skeletal Muscle Mass; PBF: percent body fat; SLM: Soft Lean Mass; FFM: Fat Free Mass; SMRA: segmental muscle right arm; SMLA: segmental muscle left arm; SMTR: segmental muscle trunk; SMRL: segmental muscle right leg; SMLL: segmental muscle left leg; BCM: body cell mass; TBM/FFM: total body mass/fat free mass

## DISCUSSION

Traditionally, an INC protocol is administered to obtain a physiological quantification (e.g. V’O_2max_) reflecting the cardiopulmonary fitness of an individual. Previous research showed that VO_2max_ is poorly correlated with the athletic performance on the court [14]. One reason seems to be the different exercise patterns between the laboratory testing procedure (i.e. INC) and the field performance of several sportive ball games (e.g. badminton, basketball, soccer). This descriptive study aimed to investigate the HIIT performance under the same relative intensity and duty cycle (120% speed and the same incline as one’s INC in the final stage alternating with a 1-min rest) on a group of 47 participants in order to explore the physiologic factors determining tolerance to effort during a HIIT. Our results showed that total number of bouts to limitation (B_lim_) was widely distributed among study population, ranging from 2.6 to 12 bouts (mean: 6.0 ± 2.6). Further regression analyses, seeking to reveal which physiological parameters determine this large distribution, suggests that HR and VO_2_ recovery are the key variables to reach a longer HIIT duration (i.e. higher number of bouts completed).

### Parameters derived from INC influencing B_***lim***_

We were interested in determining what variables extracted from a comprehensive analyses of the traditional INC test, influence B_lim_ and therefore the individual tolerance of high-intensity intermittent exercise. Results from linear stepwise regression analysis revealed that B_lim_ was more strongly associated with three physiological parameters: ΔVO_2_2, ΔHR2/, and TSI_peak_.

VO_2_ recovery (△VO_2_2), rather than VO_2max_, was found the strongest determinant of B_lim_ in the linear stepwise regression analysis. This result is in agreement with a previous study by Harris and colleagues [15] that investigated the time course of phosphocreatine (PCr) resynthesis in a group of adults undergoing a maximal test performed on cycle-ergometer. These authors observed that muscle recovery kinetics after exhaustive maximal exercise is biphasic, with alactacid component (from 10 s to a few minutes) and lactacid component (a few minutes to hours) [16, 17]. The alactacid component consists of oxygen-dependent adenosine triphosphate (ATP) and PCr replenish, which is the primary energy replenish pathway during HIIT. Participants with faster ATP/PCr replenish have more rapid VO_2_ recovery within the first minutes following the dynamic exhaustive exercise and thereby, a broader VO_2_ reserve in the next HIIT bout. Thus, as evidenced by our statistical model, these individuals are characterized by having a greater B_lim_.

The second parameter that determines performance capacity during an HIIT is the HR recovery(ΔHR2/). HR recovery consists of rapid (parasympathetic reactivation) and subsequent phase (sympathetic withdrawal and humoral factors) [18]. It reflects the regulatory capability of cardiac autonomic nervous system. Previous studies showed that HR recovery reflects the VO_2max_ [19-21] and training status [22, 23]. The present study further found that it is related to performance in HIIT. During HIIT, faster HR recovery suggests a broader HR reserve for the next bout. This highlighted the importance of cardiac autonomic nervous system regulation in determining HIIT performance.

Together with the recovery rate for V’O_2_ and HR, TSI_peak_ was found a significant parameter in explaining individual B_lim_ and performance in HIIT among the participants enrolled in the current study. TSI represents a balance between muscular oxygen delivery and consumption [24, 25]. Change in TSI has been shown to increase after HIIT training, which is accompanied by increase in mitochondrial biogenesis, capillarization, and mitochondrial enzyme activity [26-28]. It is likely that participants with lower TSI_peak_ (larger TSI change) tolerate regional metabolic acidosis better and are, thereby, prone to have greater B_lim_.

In addition, B_lim_ and age showed strong relationship in our cohort (age range 15 to 28 years) suggesting that the capacity of sustaining the HIIT for long duration is dependent by the maturity (age) of the individual undergoing the test. In a study by Ratel et al. [29], authors compared the recovery capacity among prepubescent boys (n=11, age 9.6 ± 0.7 years), pubescent boys (n=9, age 15 ± 0.7 years), and men (n=10, age 20.4 ± 0.8 years) undergoing a ten-bout intermittent sprinting cycling test (friction load = 50% optimal force), separated by 30 s, 1 and 5 min passive recovery. The capacity of maintaining peak cycling power from the first to tenth set decreased (p<0.01) by 11.3% in men and 15.3% in pubescent boy, with no changes in the prepubescent group, when the recovery interval was of 1 min, i.e. same duration we applied in the current study HIIT test. Ratel study suggested that for pubescent and men categories a longer time of recovery is needed to the higher muscle glycolytic activity and slower PCr resynthesis. Similarly, in a study by Zafeiridis et al. [30]. the effect of age was investigated respect the recovery capacity after high-intensity intermittent isokinetic strength exercise. A group of boys (age, 11.4 ± 0.5 years), teens (age, 14.7 ± 0.4 years), and men (age, 24.1 ± 2 years) were enrolled and performed two sets of exercise of 30 and 60 s bout duration, intermitted by a 1 or 2 min rest, respectively. Results showed that the teens tended to recover faster than men, suggesting that the rate of recovery for both type of task was age-dependent. Accordingly, recovery capacity from an anaerobic performance decreases with age, starting as early as 9 to 11 years. Age-related exercise capacity decline is multi-factorial, such as decrease of intramuscular PCr concentration, intramuscular creatine kinase concentration, rate of PCr hydrolysis, and glycolytic enzyme activity, as well as change in muscle architecture and speed of neural activation [16, 31-35]. However, due to the wide age range and exercise type (aerobic or anaerobic) [36-38], the detailed mechanism of age-related decline of anaerobic performance from teens to young-adult remains uncertain.

Finally, we found also that the percentage of body fat is inversely associated with B_lim_ in our cohort (r = −0.371). Higher body fat proportion means less muscle mass that can produce power. Body fat, an excessive loading to carry during any type of physical effort, has been proven to decrease the performance of anaerobic exercise [39].

### Limitations

This study has few limitations. First, the explanatory power in model 3 of the multiple linear regression model was 0.415. As consequence, other factors should be considered to explain the 1–2 bout deviation of participants’ maximal B_lim_ we seen in our cohort. A plausible factor to consider in describing which are the determinants of HIIT performance could be found on the role played by motivation and compliance of participants [14]. To minimize the variability of this latter, before data collection, each participant was informed on the relevance of the results to planning the training season. We are quite confident that view the fact that the athletes were all nationals, our motivation point was effective in obtaining reliable effort to intolerance.

In addition, it could seem contradictory that from our statistical model analyses that the recovery of HR and VO_2_ at the second minute (ΔVO_2_2, △HR2/), rather than those at 30 s or 1 min or t_1/2_, were more related to B_lim_. It can be explained by the difference between active and passive recovery. During the recovery phase, all participants were instructed to walk at their casual speed for 1 min (active recovery) and then stand still for another 2 min (passive recovery). However, HR decreased less in active recovery compared with passive recovery, in that the latter has a lower central command from the motor cortex and muscle mechanometabo receptor activity from skeletal muscle contraction than the former [40]. Oxygen consumption is also higher in active recovery [41]. The competition of oxygen between PCr replenish and muscle activity during active recovery produces higher VO_2_ [42, 43]. Due to variable casual speed during active recovery among every subject, the recovery kinetics is not comparable.

Furthermore, limitations exist in the use of NIRS methodology in estimating muscle metabolism, i.e. high melanin and large adipose tissue thickness (ATT) cause a signal attenuation, reducing the amount of light reaching the muscle tissue under investigation. Nevertheless, the low content of melanin (participants were all Asians) and the ATT below 1.8 mm confirm the reliability of our NIRS data [44]. Finally, the subjects in our study were all female. As consequence, whether sex might influence the results of our analyses deserve future investigation.

## CONCLUSION

Bouts to limitation (i.e. B_lim_) in HIIT differ greatly among individuals, provided the same relative intensity and duty-recovery cycle. VO_2_ recovery, HR recovery, and TSI_peak_ are the major determining physical factors to B_lim_. In addition, our results showed that age seems to represent a strong influencing factor. B_lim_ declines remarkably since as early as late adolescent in the study participants. The study adds interpretational value in the recovery phase of maximal incremental cardiopulmonary test and further improves our understanding of the factors that predict a good performance in HIIT.

## Abbreviations

ATT: adipose tissue thickness
B_lim_: number of total bouts to limitation (i.e. exhaustion) in high-intensity interval exercise testing
BCM: body cell mass
CWR: constant work rate testing
EqCO_2_: ventilatory equivalent for CO_2_
EqO_2_: ventilatory equivalent for O_2_
FFM: fat free mass
HHb: deoxy-(hemoglobin+myoglobin)
HR: heart rate
HR_max_: maximal HR
HR reserve: heart rate reserve
△HR0.5: HR_max_ – HR at 0.5 min recovery
△HR1: HR_max_ – HR at 1 min recovery
△HR2: HR_max_ – HR at 2 min recovery
△HR0.5/: △HR0.5 / HR_max_
△HR1/: △HR1 / HR_max_
△HR2/: △HR2 / HR_max_
HRrt_1/2_: time span the heart rate recovers to half
HIIT: high-intensity interval testing
INC: maximal incremental exercise test
MA: moderately-active subjects
Max O_2_ pulse: maximal VO_2_/HR
NIRS: near-infrared spectroscopy
O_2_Hb: oxy-(hemoglobin+myoglobin)
△O_2_Hb0.5: O_2_Hb at 0.5 min during recovery – peak O_2_Hb
△O_2_Hb1: O_2_Hb at 1 min during recovery – peak O_2_Hb
△O2Hb2: O_2_Hb at 2 min during recovery – peak O_2_Hb
△O_2_Hb0.5/: △O_2_Hb0.5/ (maximal O_2_Hb – minimal O_2_Hb) during recovery
△O_2_Hb1/: △O_2_Hb1/ (maximal O_2_Hb – minimal O_2_Hb) during recovery
△O_2_Hb2/: △O_2_Hb2/ (maximal O_2_Hb – minimal O_2_Hb) during recovery
PBF: percent body fat
SLM: soft lean mass
SMM: skeletal muscle mass
SMLA: segmental muscle mass, left arm
SMLL: segmental muscle mass, left leg
SMRA: segmental muscle mass, right arm
SMRL: segmental muscle mass, right leg
SMTR: segmental muscle mass, trunk
TBM/FFM: total body mass/fat free mass
T_1/2_: the time span the variable changes in half
THb: total-(hemoglobin+myoglobin)
TSI: tissue saturation index
TSI_peak_: nadir of tissue saturation index during INC
△TSI0.5: TSI at 0.5 min during INC recovery – peak TSI
△TSI1: TSI at 1 min during INC recovery – peak TSI
△TSI2: TSI at 2 min during INC recovery – peak TSI
△TSI0.5/: △TSI0.5/ (maximal TSI – minimal TSI) during recovery
△TSI1 /: △TSI1/ (maximal TSI – minimal TSI) during recovery
△TSI2/: △TSI2/ (maximal TSI – minimal TSI) during recovery
VCO_2_: carbonic dioxide consumption
V_E_: minute ventilation
VO_2_: oxygen consumption
VO_2max_: maximal VO_2_
△VO_2_0.5: VO_2max_ – VO_2_ at 0.5 min recovery
△VO_2_1: VO_2max_ – VO_2_ at 1 min recovery
△VO_2_2: VO_2max_ – VO_2_ at 2 min recovery
△VO_2_0.5/: △VO_2_0.5 / maximal VO_2_
△VO_2_1/: △VO_2_1 / maximal VO_2_
△VO_2_2/: △VO_2_2 / maximal VO_2_
VO_2_rt_1/2_: time span the VO_2_ recovers to half

## Acknowledgement

Authors thank all participants for the time and dedication shown to complete the experimental protocol. This research was supported by Chang Gung Memorial Hospital (grant No. CMRPG3E1561/2) and Healthy Aging Research Center, Chang Gung University and the Taiwan Ministry of Education’s Higher Education Deep Plowing Program (Grant Numbers EMRPD1H0351 and EMRPD1H0551).

## Author contribution statement

HSC, AA, and LYC conceived and designed the experiments. CSC, LHC, LYC, CCPC, and HSC performed the experiments. AA, LHC, LYC, CCPC, HCC and HSC analyzed the data. CSC, LHC, HCC and HSC contributed analysis tools. AA, CSC, LHC, and HSC wrote the paper. All authors revised, proof-read and approved the final version of the manuscript. HSC, CCPC, HCC and LYC obtained funding.

